# Preconditioning of *Caenorhabditis elegans* to Anoxic Insult by Inactivation of Cholinergic, GABAergic, and Muscle Activity

**DOI:** 10.1101/2020.08.28.266890

**Authors:** Heather L. Bennett, Patrick D. McClanahan, Christopher Fang-Yen, Robert G. Kalb

## Abstract

For most metazoans, oxygen deprivation leads to cell dysfunction and if severe, death. Sublethal stress prior to a hypoxic or anoxic insult (“preconditioning”) can protect cells from subsequent oxygen deprivation. The molecular mechanisms by which sublethal stress can buffer against a subsequent toxic insult and the role of the nervous system in the response are not well understood. We studied the role of neuronal activity preconditioning to oxygen deprivation in *C. elegans*. Animals expressing the histamine gated chloride channels (HisCl1) in select cell populations were used to temporally and spatially inactivate the nervous system or tissue prior to an anoxic insult. We find that inactivation of the nervous system for 3 hours prior to the insult confers resistance to a 48-hour anoxic insult in 4^th^-stage larval animals. Experiments show that this resistance can be attributed to loss of activity in cholinergic and GABAergic neurons as well as in body wall muscles. These observations indicate that the nervous system activity can mediate the organism’s response to anoxia.

## 1 Introduction

The function of all cells requires the constant provision of fuel and (for aerobic life), oxygen. Highly prevalent human conditions such as stroke and myocardial infarction result from a mismatch between fuel and oxygen delivery and tissue demands. During development, hypoxic insults have devastating effects on newborn infants, leading to global tissue dysfunction and disabilities.^1^ Cells can survive and adapt to low-oxygen (hypoxia) or zero oxygen (anoxia) conditions for a limited time based on cell-type specific factors and the duration or degree of oxygen deprivation. Reperfusion, the process of re-establishing blood flow to ischemic tissues, has been the principal clinical method to minimize cellular damage. However, reperfusion itself can contribute to cellular damage, thus there is an unmet need to develop better therapeutic options.^2^

A transient, sub-lethal experience of hypoxia or ischemia can protect against an otherwise lethal subsequent hypoxic-ischemic insult. This phenomenon, referred to ischemic or hypoxic preconditioning,^3–7^ suggests that cells have a latent adaptive capacity to combat the noxious effects of ischemia and hypoxia. If we understood the biochemical basis for the preconditional effect, it could potentially be harnessed for therapeutic purposes. Previous work in mammalian systems on ischemic preconditioning has highlighted a role for signal transduction pathways, (*i.e*., PI3K-AKTand ERK pathways), as well as hypoxia inducing factor (HIF).^8–10^ Nonetheless, a complete understanding of the phenomenon is lacking.

Genetically tractable organisms have proven to be a powerful platform for discovery of novel genes and pathways of biological significance. The nematode *C. elegans* prefers oxygen between 5 and 12% ^11^ and has the ability to sense and respond to shifts in oxygen that fall outside the preferred range.^12,13^ *C. elegans* are capable of surviving low oxygen stress and use a variety of pathways to achieve this depending on the degree and duration of oxygen deprivation.^14–17^ We found that ablation of the oxygen-sensing BAG (but not the URX) neurons rendered animals resistant to an anoxic insult.^13^ We postulated the neural circuit in which the BAG neuron was embedded secreted a factor(s) that heightened peripheral tissue sensitivity to anoxia. This hypothesis was derived from the observations that: 1) inhibition of neuropeptide processing (*e.g*., *C. elegans* with *egl-3* or *egl-21* mutation) and secretion (*e.g*., *C. elegans* with *unc-31* mutation) protected against anoxia, 2) nervous systemspecific rescue of *egl-3* expression in *egl-3* mutant animals restored sensitivity to anoxia, and 3) we identified one neuropeptide *nlp-40* and its receptor *aex-2* that are likely to be involved in this process.^18^ These observations highlight the cell non-autonomous determinants of *C. elegans* survival upon anoxia insult.

*C. elegans* display the preconditioning phenomenon and genetic studies implicate classical stress response pathways, genes required for lifespan, energy homeostasis, dauer formation, and genes involved cell death pathways.^15,17,19–21^ Given this starting point, we wondered if the preconditioning phenomena in *C. elegans* similarly displayed cell non-autonomous features and involved the nervous system. We designed experiments to address the following questions: 1) What is the role of the nervous system in sensing and responding to anoxic stress? And 2) What are the tissues or neuronal populations underlying the preconditioning response to anoxic stress? Our results support a role for inactivity of cholinergic and GABAergic neurons, and muscle in modulating survival to a subsequent anoxic insult.

## 2 Material and Methods

### 2.1 Strains

The following strains were used in this work. N2, referred to as wild type, CX14373 *kyEx457l [pNp403 (tag-168:HisCl1::SL2::GFP;myo-3::mCherry]* from Pokala *et al*., 2014.^22^ CX14845 *kyEx5104[pNP424 (mec-3::HisCl1::SL2::mCherry; unc-122::GFP)]*, from Pokala *et al*., 2014.^22^ CX15341 *kyEx5161[pNP488 (unc-4::HisCl1::SL2::mCherry; elt-2::mCherry)]*, from Pokala *et al*., 2014. ^22^ CX15457 *kyIs620[pNP472 (inx-1::HisCl1::SL2:GFP;myo-3::mCherry)]*, from Pokala *et al*., 2014.^22^ PS6963 *syIs336 [pHW383(Pmyo-3::nls::GAL4SK::VP64::unc-54 3’UTR; Pmyo-2::nls::mCherry)]*, from Wang *et al*., 2017.^23^ PS7160 *syIs393 [pHW504(Punc-47::nls::GAL4::VP64::let-858 3’UTR; Punc-122::RFP)]*, from Wang *et al*., 2017.^23^ PS7199 *syIs371 [pJL046(15xUAS:: Δpes-10::HisCl1::SL2::GFP::let-858 3’UTR;Punc-122::GFP)]*, from Wang *et al*., 2017.^23^ RK200 (*Punc-47::nls::GAL4SK:: VP64::let-858 3’UTR;15xUAS::Δpes-10::HisCl1::SL2::GFP::let-8583’UTR;Punc-122::GFP*), RK201[(*Pmyo-3::nls::GAL3SK::VP64::unc-54 3’UTR;15xUAS:: Δpes-10::HisCl1::SL2::GFP::let-858 3’UTR;Punc-122::GFP)]*, RK206 *sdEx5[pJP673(Punc-17::HisCl1;myo-2::mCherry)], RK207 sdEx6[pJP673(Punc-17::HisCl1;myo-2::mCherry)]*, RK210 *sdEx7[(Ptph-1::HisCl1; Pmyo-2::mCherry]*, RK211 *sdEx8[(Ptph-1::HisCl1;Pmyo-2::mCherry)]*, RK222 *sdEx9 [pJL033(Peat4::nls::GAL4SK::VP64::unc-543’UTR;Pmyo-2::nls::mCherry*, RK223 *sdEx10[pJL033(Peat-4::nls::GAL4SK:: VP64::unc-54 3’UTR;Pmyo-2::nls::mCherry)]*, RK225 *sdEx11[pJL063(Pcat-2::nls::GAL4SK::VP64::unc-54 3’UTR;Pmyo-2::nls::mCherry)]*, RK228 *sdEx12[pJL063(Pcat-2::nls::GAL4SK:: VP64::unc-54 3’UTR;Pmyo-2::nls::mCherry)]*, RK229 *sdEx13[pJL033(Peat-4::nls::GAL4SK::VP64::unc-543’UTR;Pmyo-2::nls::mCherry;3’UTR;15xUAS::Δpes-10::HisCl1::SL2::GFP::let-8583’UTR;Punc-122::GFP)]*, RK230 *sdEX14[pJL033(Peat-4::nls::GAL4SK::VP64::unc-543’UTR;Pmyo-2::nls::mCherry;3’UTR;15xUAS::Δpes-10::HisCl1::SL2::GFP::let-8583’UTR;Punc-122::GFP)]*, RK231 *sdEx15[pJL063(Pcat-2::nls::GAL4SK::VP64::unc-54 3’UTR;Pmyo-2::nls::mCherry;15xUAS::Δpes-10::HisCl1::SL2::GFP::let-8583’UTR;Punc-122::GFP)]* RK240 *sdEx16[pJL063(Pcat-2::nls::GAL4SK::VP64::unc-54 3’UTR;Pmyo-2::nls::mCherry;15xUAS::Δpes-10::HisCl1::SL2::GFP::let-8583’UTR;Punc-122::GFP)]*. All GAL4 UAS strains and plasmids were a kind gift provided by the Sternberg lab.

### 2.2 *C. elegans* culture and media preparation

Strains were reared on NGM plates seeded with *Escherichia coli* OP50 as a food source under standard conditions. NGM-H+ refers to NGM plates with 10mM Histamine dichloride added to the agar and seeded with OP50 *E. coli*. NGM-H-refers to standard NGM plates with OP50 and no histamine. Plates were prepared as described in Pokala *et al*., 2014.^22^ Synchronous populations of nematodes were generated using a 1:1 mixture of 1N NaOH and hypochlorite bleach solution for no more than 10 minutes with gravid adults. Two days later L4 stage animals were collected and assayed for survival after anoxic exposure. Protocol and details described by Theresa Stiernagle in Wormbook chapter entitled Maintenance of *C. elegans* and used in previous studies to synchronize *C. elegans* for hypoxia assays in Flibotte *et al*., 2014 and Doshi *et al*., 2019.^13,18,24^

### 2.3 Anoxia exposure and assessment of survival

All experiments were performed on L4 stage animals. For anoxic insult, 30 mid L4 stage animals per genotype were picked to an NGM plate, which was placed in a Bio-Bag (Type A anaerobic environmental system, Becton-Dickinson Company, Franklin Lakes, New Jersey), anoxic conditions were induced and maintained for 48 hours at 20°C as described previously.^13,25^ Bags were then opened, and animals were allowed to recover in ambient oxygen for 24 hours before being scored for survival. Surviving animals were identified as those that moved spontaneously or after gentle prodding with a platinum wire. Most resumed feeding and matured into egg-laying adults.

### 2.4 Preconditioning paradigm with histamine

Early L4 stage animals, as judged by vulval morphology, were plated on NGM-H+ or NGM-H-plates for either 0.5, 1 or 3.5 hours. Animals were then moved to NGM-H-plates for 1.5 hours and subsequently exposed to anoxic conditions.

### 2.5 Preconditioning paradigm to starvation

Animals were fed OP50 *E. coli* until the early L4 stage of development. Early L4 stage animals were transferred to either NGM plates seeded with OP50 *E. coli* or unseeded plates for a period of 3.5 hours. Experimental animals were taken off NGM unseeded plates and placed onto NGM plates containing OP50 for 1.5 hours. Control animals were always exposed to conditions where food was plentiful.

### 2.6 Activity Assays

The “WorMotel” device, a multi-well imaging platform, was used to assay *C. elegans* activity for individual animals for a 3.5 hour period.^26^ Device construction, *C. elegans* cultivation, and imaging setup were performed as described in Churgin and Fang-Yen 2017 ^26^ except images were recorded every 10 seconds.

### 2.7 Statistical Analysis

All statistical analysis and graph construction were prepared using GraphPad Prism version 8 (San Diego, California) or MATLAB (Natick, Massachusetts). We averaged survival results from 3+ independent trials performed on different days. Experiments are done in triplicate with 30 animals per genotype or condition. Error bars indicate the standard error of the mean for all experiments. Significant differences were assessed by paired Student’s t-tests (two tailed) for differences between two groups. For groups of three or more, the survival was analyzed by one-way ANOVA, followed by a Tukey’s multiple comparisons post hoc test. Significance was considered if p< 0.05.Video recordings of *C. elegans* behavior were analyzed using a MATLAB script as previously described in Churgin and Fang-Yen 2017.^26^ The relationship between statistical power and effect size (Figure 4b) was determined using MATLAB and assuming normally distributed data with variance equal to that observed in the real activity data. The Anderson-Darling test was used prior to statistical testing to determine whether data were consistent with a normal distribution.

**Figure 1:**
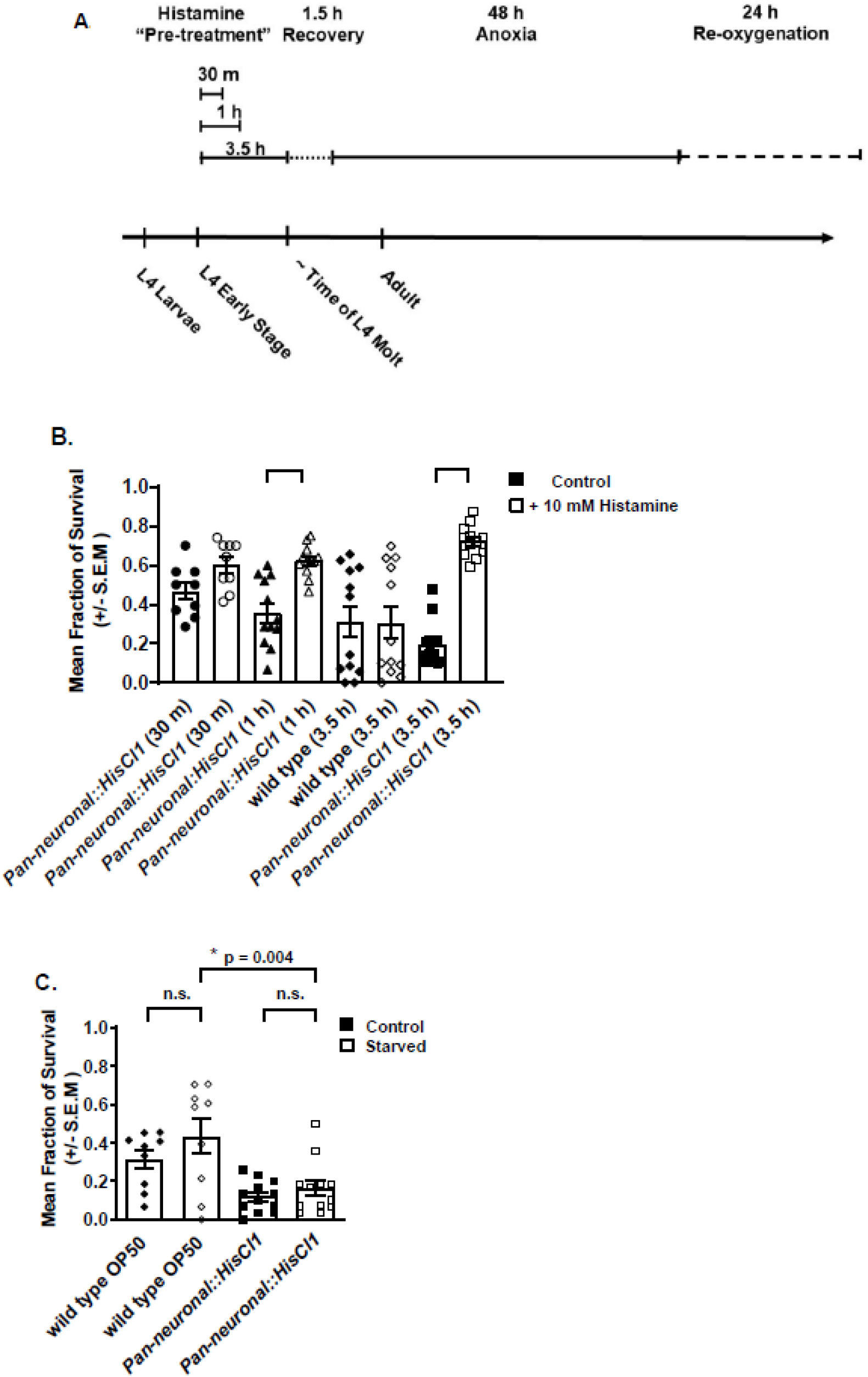
Impaired Neuronal Activity Prior to Anoxic Insult Increases Survival. **A.) Preconditioning to 48 hours of anoxia experimental paradigm.** Thirty animals carrying the histamine gated chloride channel behind a neuronal or a tissue specific promoter were selected as early L4 animals. Animals were placed on NGM plates containing either 10mM of histamine seeded with OP50 *E. coli*for 30 minutes, 1 hour, or 3.5hours. Control animals placed to NGM OP50 *E. coli* plates lacking histamine. Animals recovered on non-histamine plates for 1.5 hours and then asphyxiated for 48 hours. Fraction of survival was scored for animals that developed into adults, regained movement and resumed feeding 24 hours after anoxic insult. **B.) Inactivation of the nervous system, prior anoxic insult has a beneficial effect.** Wild type animals and animals carrying the histamine gated chloride channel 1 behind a pan-neuronal *tag-168* promoter were treated as described in panel A. Controls (black filled shapes) and experimental histamine exposed (non-filled shapes). Results are shown for 4 independent trials, n=360 animals per condition **C.) Starvation prior to 48 hours of anoxia has no impact on survival.** Wild type and animals expressing histamine gated chloride channel were selected as early L4 animals to NGM plates seeded with OP50 *E. coli* (control black filled shapes) or plates with no OP50 *E. coli* (starved non-filled shapes). Results are shown for 3 independent trials (n=270 animals) wild type and 4 independent trials pan-neuronal histamine strain, n=360 animals per condition.

**Figure 2:**
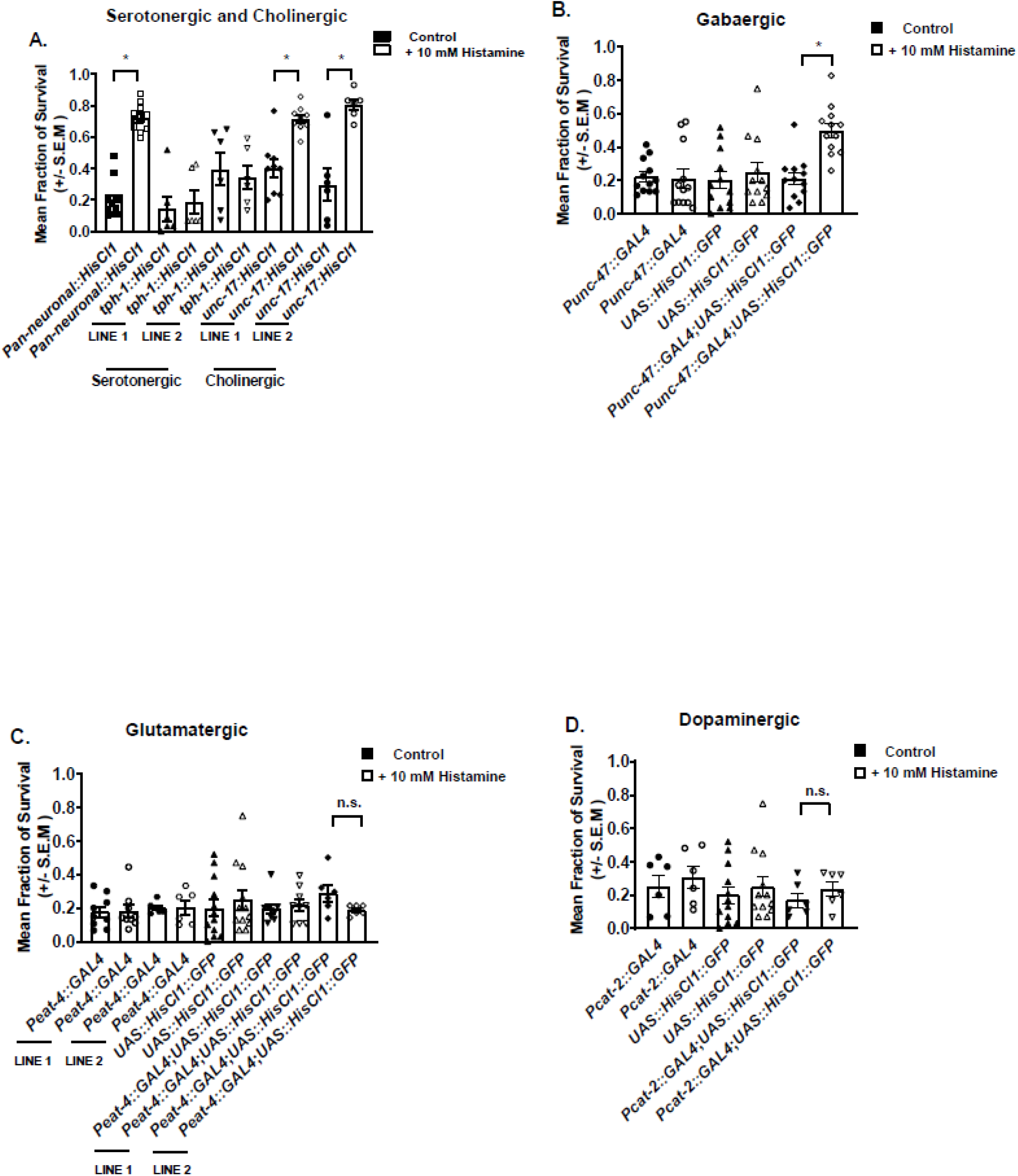
Inactivity of Cholinergic and GABAergic Neurons Mediates the Preconditioning Effect. **A.) Inactivation of cholinergic neurons, prior to anoxic insult has a beneficial effect.** Wild type animals and animals carrying the histamine gated chloride channel under the pan-neuronal promoter *tag-103*, cholinergic promoter, *unc-17*, or serotonergic promoter *tph-1* were selected to control plates (black filled shapes) or 10mM histamine plates (non-filled shapes) as early L4 animals, prior to anoxic stress. * denotes significance. Results are shown for 4 independent trials, n=360 animals per condition. **B.) Loss of GABAergic signaling prior to anoxic insult confers a survival benefit.** Wild type, animals expressing the GABAergic promoter (*Punc-47*) behind the GAL4 sequence, animals carrying the histamine gated chloride channel behind the UAS activated sequence, as well as animals expressing GAL4 under the GABAergic promoter with the UAS histamine gated chloride channel were tested. * denotes significance *Punc-47::GAL4:15xUAS::HisCl1::SL2::GFP* control versus experimental. **C.) Loss of glutamatergic signaling does not precondition animals to anoxia.** Wild type animals, animals expressing the glutamatergic promoter (*Peat-4*) behind the GAL4 sequence, animals carrying the histamine gated chloride channel behind the UAS activated sequence, as well as animals expressing GAL4 under the *eat-4* promoter with the UAS histamine gated chloride channel were tested as in panel B. Results are shown for 2 independent trials. **D.) Inactivation of dopaminergic pathway prior to anoxia does not yield a survival advantage.** Wild type animals, animals expressing the dopaminergic promoter (*Pcat-2*) behind the GAL4 sequence, animals carrying the histamine gated chloride channel behind the UAS activated sequence, as well as animals expressing GAL4 under the *cat-2* promoter with the UAS histamine gated chloride channel were tested as in panel B. Results are shown for 2 independent trials.

**Figure 3:**
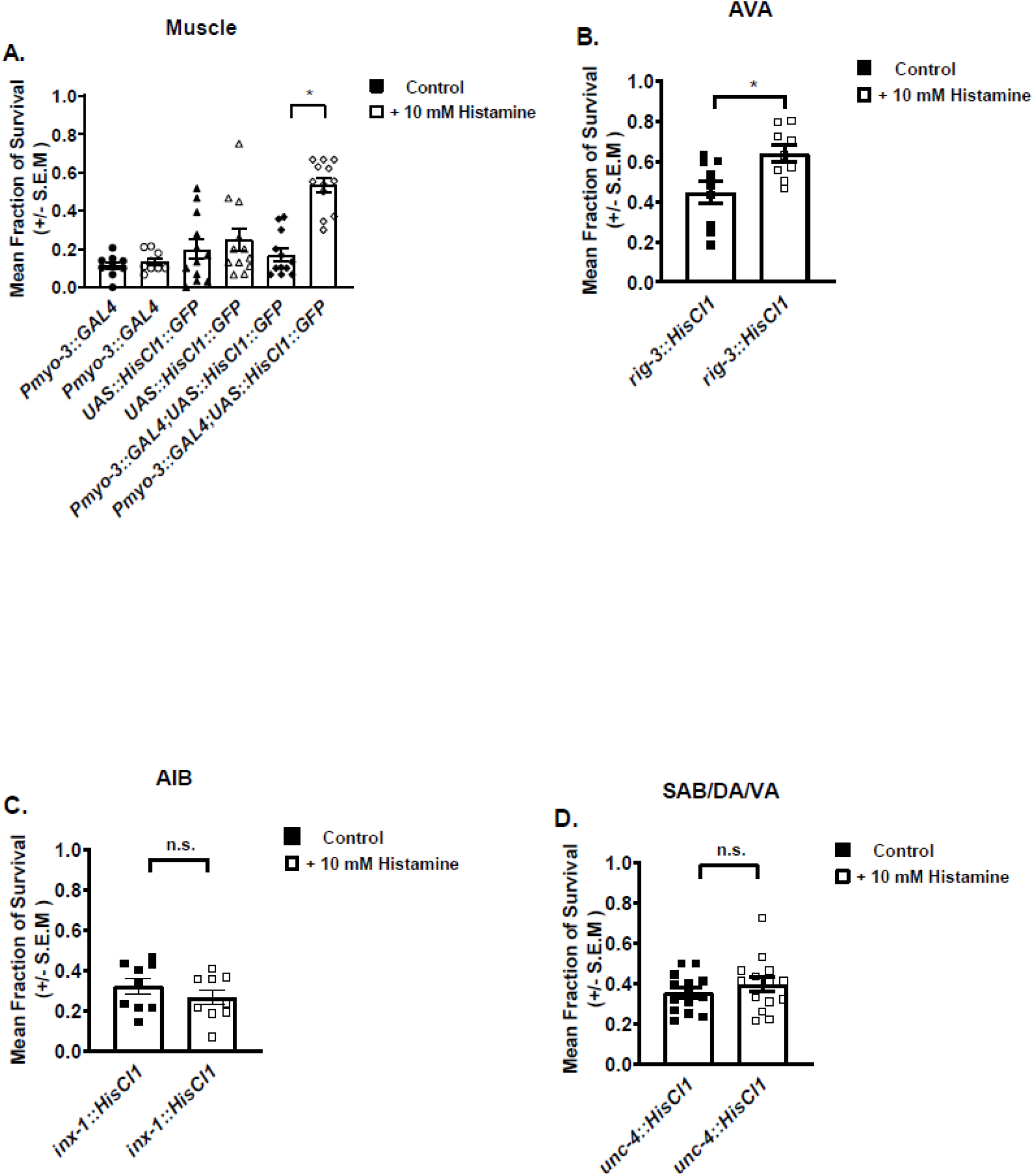
Neuromuscular Activity Mediates the Preconditioning Effect to Anoxia. **A.) Inactivation of the muscle prior to anoxic insult has an advantageous effect on survival.** Wild type, animals expressing the muscle promoter (*Pmyo-3*) behind the GAL4 sequence, animals carrying the histamine gated chloride channel behind the UAS activated sequence, as well as animals expressing GAL4 under the muscle promoter with the UAS histamine gated chloride channel were selected to 10mM histamine or control plates as early L4 animals 3.5 hours, prior to anoxic stress. Fraction of survival was scored after 24 hours. Significance between *Pmyo-3::GAL4:15xUAS::HisCl1::SL2::GFP* control versus experimental; error bars respresent the SEM. Results are shown for 4 independent trials. **B.) Inactivity of command interneurons AVA yields a survival advantage to anoxia.** Animals carrying the histamine gated chloride channel behind the *rig-3* promoter were selected as early L4 animals. Controls animals (black filled shapes) and animals exposed to 10mM histamine, experimental condition, (non-filled shapes) as early L4 animals, prior to 48 hours of anoxic stress. Fraction of survival was scored after 24 hours. Results are shown for 4 independent trials. **C.) Inactivity of AIB interneuron prior to anoxic insult does not provide a survival advantage.** Animals expressing the histamine gated chloride channel behind the *inx-1* promoter were tested, experimental paradigm as in panel C. Results are shown for 4 independent trials. **D.) SAB-DA-VA motor neuron inactivity is dispensable for the preconditioning response to anoxic insult.** Animals expressing the histamine gated chloride channel behind the *unc-4* promoter were tested for pre-conditioning response to 48hours of anoxic insult experimental paradigm as in panel C. Results are shown for 4 independent trials.

**Figure 4:**
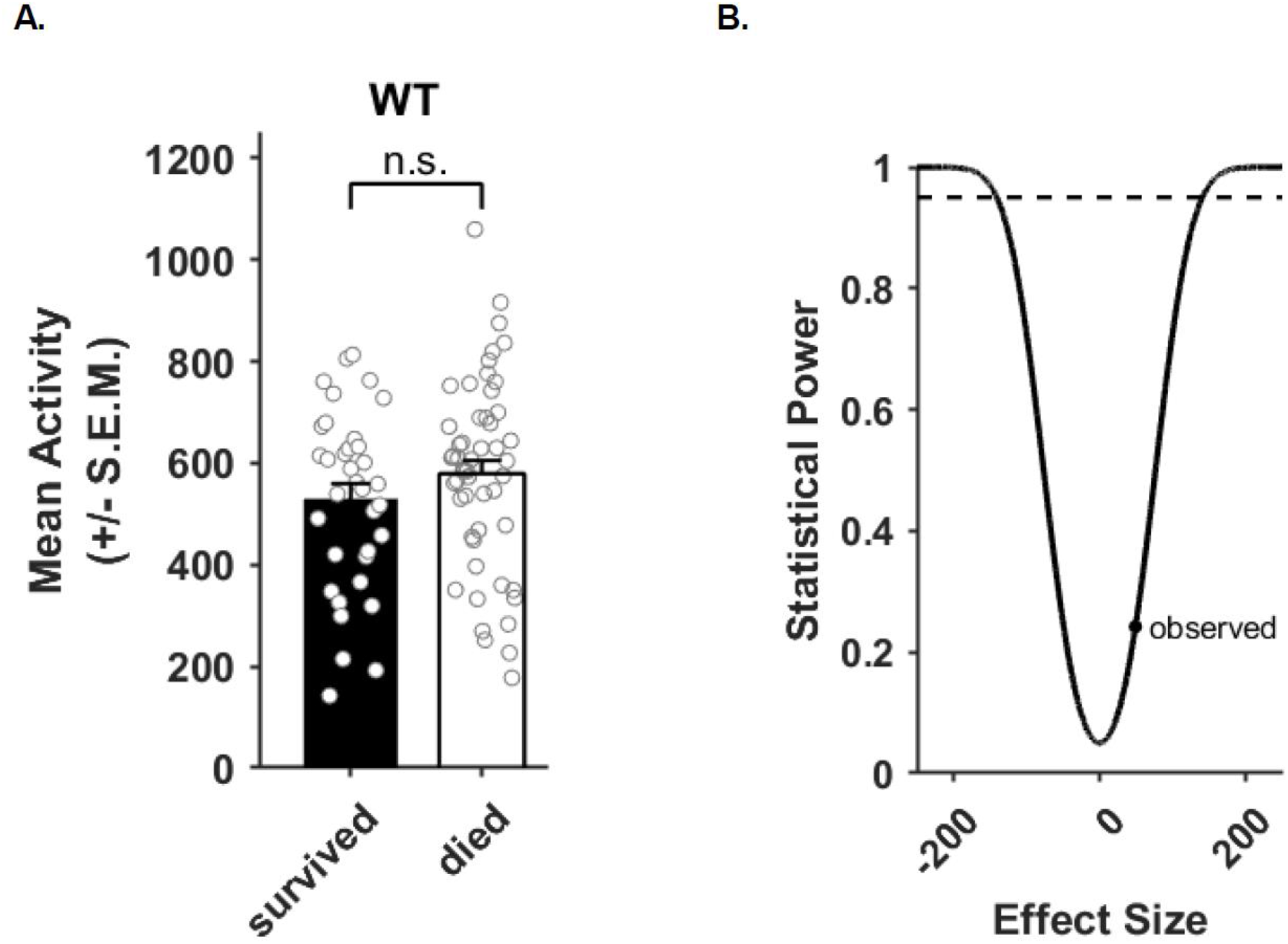
Survival is Not Related to Locomotor Activity Prior to Anoxic Insult. **A.) Activity of survivors and non-survivors prior to anoxic insult.** Wild type animals were selected at the early L4 stage and loaded into the “WorMotel” device and assayed for locomotor activity for a 3.5 hour period. Dots indicate activity of individual animals. Results are shown for 87 individual animals (35 survivors and 52 non-survivors) from five replicates. **B.) Statistical power analysis.** The statistical power (probability of rejecting the null hypothesis of no difference in activity between survivors and non-survivors if the null hypothesis were not true) of our experiment for different effect sized based on the variance and sample sizes in panel A is shown (black line) A statistical power of 0.95 is indicated by a horizontal dashed line, and the observed effect size from panel a is indicated by a dot.

### 2.8 Data Availability Statement

The data that support the findings of this study, all primary data, including number of animals scored, and number of independent replicates is available from the corresponding author upon reasonable request.

## 3 Results

### 3.1 Hyperpolarization of the Nervous System Prior to Anoxic Insult Yields a Survival Benefit

We hypothesized that an animal’s susceptibility to an anoxic insult would be influenced by nervous system activity preceding the insult. To test this idea, we used a chemo-genetic approach based on the transgenic expression of histamine gated chloride channels (HisCl1) in select populations of cells. Wild type *C. elegans* neither express HisCl1 nor synthesize histamine. Exogenous provision of histamine to transgenic *C. elegans* expressing HisCl1 in neurons leads to hyperpolarization and reduced activity.^22^

We began by studying animals expressing the HisCl1 expressed throughout the nervous system via the *tag-168* promotor (*i.e*., pan neuronal (pn) HisCl1s). In the absence of histamine, the animals with pnHisCl1 appeared and behaved like wild type animals, as previously reported.^22^ When animals with pnHisCl1 expression were placed on nematode growth media agar plates supplemented with histamine (NGM-H+) they became paralyzed in about 2 minutes.

Next we asked if a brief period of paralysis prior to anoxia influenced survival after an anoxic insult. To study this, early L4 stage pnHisCl1 animals were placed on NGM-H+ for either 30 minutes,1 hour, or 3.5 hours, then transferred to standard NGM plate for 1.5 hours, where they regained locomotor ability. When then subjected to 48 hours of anoxic insult and assessed after 24 hours of normoxic recovery, we found that pnHisCl1 animals that had been exposed to histamine (NGM-H+ plates) had increased survival relative to pnHisCl1 animals (that had been grown on NGM-H-plates) (Figure 1B). There was a trend towards increased survival at 1 hour and a statistically significant beneficial effect was seen in pnHisCl1 animals with 3.5 hours of nervous system inactivity prior to anoxia (pnHisCl1 NGM-H-0.31 +/- 0.06 versus pnHisCl1 NGM-H+ 0.73 +/-0.02, ANOVA *F_(7,82)_* = 12.96, P< 0.0001, Figure 1B). As a result, this time point is used as the pre-conditioning manipulation for all other strains. Exposure of wild type animals to histamine conferred no anoxia survival benefit. (N2 NGM-H-0.31 +/- 0.08 versus N2 NGM-H+ 0.30 +/- 0.08 Figure 1B). These results indicate that inactivity of the entire nervous system prior to an anoxia insult protects against a subsequent anoxic insult (*i.e*., preconditioning).

pnHisCl1 animals, when grown on NGM-H+ plates are paralyzed and display no pharyngeal pumping (feeding) behavior. We considered the possibility that inhibition of pharyngeal pumping, which would impede food intake for 3.5 hours, might activate a stress response pathway that protects animals against anoxic stress. This is suggested by prior work showing that a period of starvation, referred to as starvation induced stress response or caloric restriction, and can lead to stress resistance and increase in longevity.^27–29^ To examine this issue, we reared wild type animals on NGM OP50 plates, and at the early L4 stage transferred animals to NGM plates with or without OP50 (starvation condition). Animals were then transferred to NGM plates with OP50 (food) for 3.5-hours and subsequently subjected to anoxia. We find that 3.5 hours of starvation prior to anoxia does not enhance survival after an anoxic insult (N2 +food 0.31 +/-0.05 versus N2 starvation 0.43 +/-0.09, pnHisCl1 +food 0.13 +/-0.02 versus pnHisCl1 starvation 0.15 +/-0.04 ANOVA, *F_(3,38)_* = 7.444, P=0.924 Figure 1C). These results show that the nervous system inactivity is unlikely to protect against anoxic injury owing to a brief period of starvation.

### 3.2 Hyperpolarization of Cholinergic and GABAergic Signaling Preconditions *C. elegans* to Anoxic Stress

To determine which neuronal population confers survival benefit when inactivated, we studied animals with HisCl1 in neurochemically defined classes of neurons. To study cholinergic or serotonergic neurons we generated transgenic animals in which the *unc-17* or *tph-1* promoter (respectively) drove expression of HisCl1. Four independent extrachromosomal array lines were generated and tested. Both groups of animals (HisCl1 in cholinergic neurons: ch-HisCl1, or in serotonergic neurons: ht-HisCl1) appeared normal on NGM-H-plates. When placed on NGM-H+ plates the ch-HisCl1 animals became paralyzed, while the ht-HisCl1 displayed no overt phenotype. In our pre-conditioning paradigm, we find that inactivation of cholinergic activity, but not serotonin activity, for 3.5 hours prior to an anoxic insult confers a survival benefit (ch-HisCl1 NGM-H-0.40, +/-0.06 and 0.30, +/- 0.10 lines 1 and 2 respectively versus ch-HisCl1 NGM-H+ 0.72, +/- 0.03 and 0.81, +/-0.03 lines 1 and 2 respectively ht-HisCl1 NGM-H- 0.20, +/-0.08 and 0.40, +/-0.10 lines 1 and 2 respectively ht-HisCl1 NGM-H+ 0.19, +/-0.07 and 0.34, +/-0. 07 lines 1 and 2 respectively ANOVA *F_(9,68)_* = 18.83 p value < 0.0001, Figure 2A).

Next, we used the bipartite GAL4-UAS system to study other neurochemically defined neuronal populations. ^23^ Animals containing UAS sequences driving HisCl1 were crossed to animals in which the *unc-47* promoter drives GAL4 to generate animals expressing the HisCl1 in GABAergic neurons (ga-HisCl1). Ga-HisCl1 animals appeared normal on NMG-H-plates, but displayed a severely uncoordinated phenotype, *i.e*. abnormal body wall contraction or defective movement when prodded with the platinum wire pick, when placed on NGM-H+ plates. In our pre-conditional paradigm, we find that loss of GABAergic activity for 3.5 hours prior to 48 hours of anoxia conferred a survival benefit (ga-HisCl1-NGM-H-0.21 +/- 0.04 versus ga-HisCl1-NGM-H+ 0.50 +/- 0.04, ANOVA *F_(5,66)_* =6.089, p value= 0.0001, Figure 2B).

To generate *C. elegans* with glutamatergic or dopaminergic expression of HisCl1, constructs containing *eat-4*, and *cat-2* promoter regions driving GAL4 sequences, along with plasmids containing the UAS sequences driving the HisCl1were injected into N2 animals to generate extrachromosomal array lines. Four independent lines were generated with each glutamatergic (glu-HisCl1) and dopaminergic (dop-HisCl1) construct. glu-HisCl1 and dop-HisCl1 animals appeared normal on standard NGM plates that lacked histamine, and displayed no overt phenotype on NGM-H+ plates. We find that loss of neither glutamatergic nor dopaminergic activity for 3.5 hours prior to anoxic insult conferred a survival benefit (glu-HisCl1 NGM-H-0.20 +/-0.04 and 0.29 +/- 0.05 lines 1 and 2 respectively versus glu-HisCl1 NGM-H+ 0.18 +/-0.04 and 0.19 +/- 0.01 lines 1 and 2 respectively, ANOVA *F_(9,74)_ =0.5165* p=0.8582. dop-HisCl1 NGM-H-0.17 +/-0.04 versus dop-HisCl1 NGM-H+ 0.23 +/-0.04 ANOVA *F_(7,46)_ =4.551* p=0.996 Figure 2C-D respectively). Combined, these results suggest that activity of cholinergic or GABAergic neurons regulate the pre-conditioning response to anoxia.

### 3.3 Hyperpolarization of Muscle Activity Preconditions *C. elegans* to Anoxia

Given that loss in cholinergic or GABAergic activity led to paralysis and impaired locomotion respectively, and conferred survival prior to anoxic stress, we asked whether muscle inactivity prior to anoxia would also yield a survival benefit. To test this, we used the UAS-GAL4 system to express HisCl1 in body wall and vulval muscle cells.^30,31^ Animals expressing HisCl1 in muscles on NGM-H-plates were indistinguishable from N2 *C. elegans*. When placed on NGM-H+ plates the muscle-HisCl1 animals became paralyzed and this effect reversed when subsequently moved to NMG-H-plates. We found that 3.5 hours of muscle paralysis prior to 48 hours anoxic insult confers a survival benefit (muscle NGM-H- 0.17, +/- 0.03 versus muscle NGM-H+ 0.53, +/- 0.03 ANOVA *F_(5,60)_ =13.79*, p value <0.0001 Figure 3A). This might indicate that active muscle secretes a factor that makes the organism sensitive to anoxia. Or inactive muscle secretes a factor that makes the organism resistant to anoxia. Regardless of the mechanism, these results suggest that reducing muscle activity below a threshold contributes to the preconditioning phenomenon.

### 3.4 The Preconditioning Response to Anoxia is Dependent on AVA Command Interneurons

To further probe the neural circuit regulating the preconditioning response to anoxia, we studied the role of select interneurons. We tested AVA command interneurons first because these neurons facilitate backward locomotion in the animal, receive acetylcholine neurotransmitter input, and make connections onto motor neurons. To determine if AVA command interneurons play a role in in mediating the preconditioning response to anoxia, we studied animals expressing HisCl1 under the control of the *rig-3* promoter (AVA-HisCl1).^32^ AVA-HisCl1 animals appeared normal on NGM-H-plates and displayed a mild uncoordinated (Unc) phenotype when placed on NGM-H+ plates. Animals with a loss in AVA activity for 3.5 hours prior to 48-hour anoxic insult had increased survival compared to controls (AVA-HisCl1 NGM-H-0.44,+/- 0.06 versus AVA-HisCl1 NGM-H+ 0.64, +/- 0.04, *t_(8)_=2.971*, p=0.01, paired t test, Figure 3B).

Next we tested a different population of interneurons that are also part of the locomotor circuit. AIB are a pair of amphid interneurons that receive input from sensory neurons, make connections onto motor neurons, and also regulate locomotion in *C. elegans*. Animals expressing HisCl1 in AIB interneuron pair on NGM-H- and NGM-H+ plates are indistinguishable from N2 *C. elegans*. In our preconditioning paradigm, we find that impairing AIB interneuron activity prior to anoxia did not yield a survival benefit (AIB-HisCl1 NGM-H 0.32 +/-0.04 versus AIB-HisCl1 NGM-H+ 0.26 +/-0.04, *t(8)=1.215*, p=0.258, paired t test Figure 3C). This suggests that the pre-conditioning phenomenon involves the activity within specific neurons of the locomotor circuit.

Since AVA interneurons make connections with motor neurons, we asked if inactivity of specific motor neurons prior to anoxia might lead to increased survival. We tested DA and VA motor neurons for several reasons: 1) they innervate dorsal and ventral muscles respectively,^33^ 2) they receive cholinergic input,^34^ 3) they receive direct input from AVA interneurons to initiate backward locomotion‘^35^ and 4) AVA hyperpolarization maybe mediated by inactivation of about 2/3 of cholinergic motor neurons, including classes DA and VA motor neurons through gap junctions.^33,36,37^ To determine if DA and VA motor neurons play a role in in mediating the preconditioning response to anoxia, we studied animals expressing HisCl1 under the control of the *unc-4* promoter, which also expresses in three SAB head motor neurons (SAB-DA-VA-HisCl1).^38^ SAB-DA-VA-HisCl1 animals appeared normal on standard NGM-H-plates and displayed a weak phenotype when placed on NGM-H+ plates. We find that inactivity of SAB, DA, and VA motor neurons in our preconditioning paradigm does not confer a survival after anoxic insult. (SAB-DA-VA NGM-H-0.35 +/-0.03 versus SAB-DA-VA NGM-H+ 0.39 +/-0.04, *t_(13)_*=1.172, p=0.262, paired t-test, Figure 3D).

### 3.5 Lack of Locomotor Activity Prior to Anoxic Insult is Not a Predictor of Survival

Our results show that hyperpolarization of certain population of neurons or muscle prior to anoxic insult either impaired or paralyzed animals, and led to a survival benefit. We asked whether the amount of locomotor activity prior to anoxia could predict survival after an anoxic insult in untreated wild-type animals. To this end we monitored spontaneous activity of N2 animals prior to 48-hours of anoxic stress using a multi-well imaging platform (WorMotel) and image analysis. Within a population of *C. elegans*, individuals display variations in locomotor activity. We compared the pre-anoxia activity of animals that survived anoxia to those that died. We found only a small, non-significant difference of 49.2 activity values between survivors and non-survivors (two tailed t-test, p value = 0.22, Figure 4A). Based on the sample size and variance of our dataset, the smallest difference in activity we could have reliably detected (statistical power = 0.95) was 140.8 activity values (Figure 4B). Collectively, these result suggests that, while normal *C. elegans* vary in their spontaneous activity levels, they are operating above the threshold that evokes the pre-conditioning phenomenon.

## 4 Discussion

The preconditioning phenomena is a physiological process that raises the threshold for cellular damage evoked by environmental insults. A mechanistic understanding of this process might be harnessed for therapeutic ends. Here we show that activity of specific set of neurons and muscle cells have a substantial impact on the susceptibility of developing nematodes to anoxic insult. Since the preconditioning phenomena has been described throughout the animal kingdom, insight into this physiological response in a genetically tractable organism may have broad application.^39–41^

One salient feature of our investigations here and in prior publications (Flibotte *et al*., 2014 and Doshi *et al*.,2019) is variability of survival after a 48-hour anoxic insult. We considered potential sources of this in our isogenic population of *C. elegans* (*i.e*., modest differences in animal age, number of animals on a plate, prior history of starvation, distance of plate from catalyst that induce anoxic conditions, number of plates in a biobag and age of NGM plates) and none appear to account for the variability. We suspect that natural stochasticity in biological systems may be the underlying source of the variability we see. This is a topic of great interest to the *C. elegans* community ^42–46^ and an area of active inquiry. Regardless of the source, by undertaking many independent trials and reporting averages we have worked to control for this variability. Using this approach, we believe we can draw valid conclusions despite the unavoidable variability.

A substantial amount of research into the preconditioning phenomena comes from investigations of heart tissue. Two temporally distinct phases in ischemic-reperfusion models have been described; early (or “first window of protection”, lasting ~ 2-3 hours) and late (“second window of protection”, onset 12-24 post preconditioning and lasting ~72-90 hours).^41,47–49^ Much of the physiology, cell biology and molecular biology is studied at the level of the heart itself – for example, transient interruption of coronary blood flow prior to a vessel occlusion reduces cardiac infarction size.^5^ Another form of cardiac preconditioning is termed “remote” because it is elicited by inducing transient ischemia of distal organs such as the small intestine, kidney and skeletal muscle.^50–53^ Humeral factors are posited to be released from extracardiac organs in this paradigm which confer stress resistance on the heart. Neuronal pathways and systemic responses may also be involved.^40,54^

Early phase cardiac precondition involves the local release of factors such as reactive oxygen or nitrogen species, bradykinin and adenosine.^41,55^ In parallel these agents activate several signaling cascades that include Akt, Erk1/2, protein kinase C^56^ and lead to the opening of mitochondrial ATP-sensitive potassium channels (K_ATP_).^19,57^ This has been termed the reperfusion injury salvage kinase (“RISK”) pathway.^55^ The cardioprotective effects are thought to be related to opposition of the mitochondrial permeability transition pore opening by active K_ATP_. Another pathway that is involved in cardio-protection (“survivor activation factor enhancement”) involves TNF- and the JAK/STAT pathway.^58^

Precisely how these cardioprotective pathways are coordinated and regulated remains an area of active investigation. Late phase cardiac preconditioning appears contingent on early phase signals and on transcription and translation. Remote preconditioning bears the signature of both early and late preconditioning (*i.e*., involvement of adenosine, bradykinin, *etc*.) but the nature of the humeral factors, the putative receptors and signaling processes are unknown.^41,55^

Several groups have studied the preconditioning phenomena in *C. elegans*. The Crowder group showed that unfolded protein response component IRE-1 (in a pathway independent XBP-1) and GCN-2 (in a pathway independent of phosphorylation of translation factor eIF2α) mediate the pre-conditioning response to hypoxia.^19,20^ In addition, they implicated the apoptosis factor CED-4 (also known as *apaf-1*) in a novel mechanism that does not require any other known core apoptosis genes.^20,59^ Genetic pathways that regulate energy dynamics have roles in the preconditioning response. The Padilla group showed that survival to anoxia was dependent on the energy sensor AMP regulated protein kinase (AMPK).^59^ This same beneficial effect could be mimicked by exposing animals to the dietary restriction-like state induced by metformin.^59^ Work from the Miller group showed that a several hours of fasting blunted protein homeostasis defects evoked by hypoxia and that this involved the insulin/insulin-like growth factor receptor *daf-2* but not its downstream target, *daf-16*^60^ Collectively, these studies provide valuable information about the genetic underpinning of the preconditioning phenomena, however it remains to be determined if these genes work in a single pathway or multiple parallel pathways. In addition, these studies do not provide insights into the cell autonomous versus cell non-autonomous contributions to the preconditioning phenomena.

What accounts for this heightened state of resistance? One interpretation is that inactivation of neuronal populations that impair movement suspend natural development in early L4 stage animals and these developmentally younger animals are inherently resistant to anoxic insult. However, we think that is unlikely because previous work showed no difference in survival to anoxic stress between early versus late stage animals.^18^ We therefore consider two, not mutually exclusive, possibilities to explain these observations. First, normal physiological activity of cholinergic and GABAergic neurons and muscle might secrete an “anoxia sensitivity factor” which heightens organismal vulnerability to anoxia. When these cells are electrically silenced, the abundance of this putative factor is reduced temporarily and thus organisms display increased rates of survival after an anoxic insult. Second, cholinergic, GABAergic neurons and muscle that are electrically silenced might secrete an “anoxia resistance factor”. This putative factor temporarily increases the resistance of the organism to an anoxia insult. These considerations are aligned with well-described cell non-autonomous stress signaling in *C. elegans*; such signals can originate from distinct populations of neurons as well as glial cells.^61–65^ A goal of future studies should be to determine whether the preconditioning phenomenon evoked by muscle inactivity (for example) is due to a sensitivity versus a resistance factor. Understanding the biochemical nature of this putative factor and how its signaling affects the response to an anoxic insult will be of enormous interest.

Finally, we note that the work described herein is unique in that the preconditioning stimulus is not an abbreviated exposure to an otherwise toxic insult. This differs from remote preconditioning wherein short duration ischemia to the heart or distal organs influences the outcome of a subsequent coronary vessel occlusion.^5^ It differs from the work of Dasgupta *et al*. in which 4 hours of hypoxia followed by a 24 hours recovery period afforded protection against 24 hours of hypoxia.^20^ We retain the nomenclature of preconditioning since it is a transient manipulation prior to a severe insult that moderates outcome, although this designation is arguable. We believe that expanding the notion of preconditioning in this way can help identify physiological states of higher or lower susceptibility to insult that are dynamic and susceptible to manipulation.

## 5 Conclusion

The role of the nervous system in preconditioning to anoxic insult has not been extensively studied. The neurons and tissues involved in modulating the preconditioning response to anoxia are not known. Our results implicate cholinergic, GABAergic, and muscle activity in mediating the preconditioning response to anoxic insult. Our observations raise several questions that should be addressed in the future. What is special about the cholinergic and GABAergic neurons (as opposed to other neurochemically defined neurons that also impact muscle function) that is particularly beneficial to *C.elegans* under standard cultivation conditions? What signals are elaborated by cholinergic and GABAergic neurons and muscle cells that heightens susceptibility of anoxia? Do all tissues respond to these signals or is there a cascade of signal transduction from tissue to tissue? Insight into these issues may bring us closer to harnessing the preconditioning phenomena for therapeutic use.

## 6 Acknowledgements

We would like to thank Cornelia Bargmann and Paul Sternberg for generously sharing with us several strains to test in our experimental paradigm. We would like to thank Michael Crowder at the University of Washington Seattle and David M. Raizen at the University of Pennsylvania for providing intellectual support and guidance in our experimental design and results. We also thank the Worm Group laboratories at the University of Pennsylvania for their feedback and support. We thank the members of the Kalb, Fang-Yen, Burkhardt, Argon, laboratories for support. We would like to thank Janis Burkhardt, Yair Argon, and Steven Seeholzer for generously providing reagents, equipment, and space to conduct our experiments. We also thank Erika Perez at Xavier University of Louisiana and Gabriel Perron at Bard College for guidance on statistical analyses. Some strains were provided by the CGC, which is funded by NIH Office of Research Infrastructure Programs (P40 OD010440).

H.L.B and R.G.K designed the experiments and wrote the manuscript. All authors proofread manuscript. H.L.B, and P.M conducted experiments and constructed figures. H.L.B analyzed all results.

The research reported in this publication was supported by the National Institutes of Health K12GM081259 (H.L.B), NS087077 and NS05225 (R.G.K.) and the Les Turner ALS Center at Northwestern University (R.G.K.).

## References

1. Volpe JJ. Perinatal brain injury: from pathogenesis to neuroprotection. Ment Retard Dev Disabil Res Rev. 2001;7(1):56–64.

2. Powers SK, Murlasits Z, Wu M, Kavazis AN. Ischemia-reperfusion-induced cardiac injury: a brief review. Med Sci Sports Exerc. 2007;39(9):1529–1536.

3. Bolli R. Preconditioning: a paradigm shift in the biology of myocardial ischemia. Am J Physiol Heart Circ Physiol. 2007;292(1):H19–27.

4. Dirnagl U, Becker K, Meisel A. Preconditioning and tolerance against cerebral ischaemia: from experimental strategies to clinical use. Lancet Neurol. 2009;8(4):398–412.

5. Murry CE, Jennings RB, Reimer KA. Preconditioning with ischemia: a delay of lethal cell injury in ischemic myocardium. Circulation. 1986;74(5):1124–1136.

6. Obrenovitch TP. Molecular physiology of preconditioning-induced brain tolerance to ischemia. Physiol Rev. 2008;88(1):211–247.

7. Pignataro G, Scorziello A, Di Renzo G, Annunziato L. Post-ischemic brain damage: effect of ischemic preconditioning and postconditioning and identification of potential candidates for stroke therapy. FEBS J. 2009;276(1):46–57.

8. Hyvarinen J, Hassinen IE, Sormunen R, et al. Hearts of hypoxia-inducible factor prolyl 4-hydroxylase-2 hypomorphic mice show protection against acute ischemia-reperfusion injury. J Biol Chem. 2010;285(18):13646–13657.

9. Liu D, He M, Yi B, Guo WH, Que AL, Zhang JX. Pim-3 protects against cardiomyocyte apoptosis in anoxia/reoxygenation injury via p38-mediated signal pathway. Int J Biochem Cell Biol. 2009;41(11):2315–2322.

10. Ravingerova T, Matejikova J, Neckar J, Andelova E, Kolar F. Differential role of PI3K/Akt pathway in the infarct size limitation and antiarrhythmic protection in the rat heart. Mol Cell Biochem. 2007;297(1-2):111–120.

11. Gray JM, Karow DS, Lu H, et al. Oxygen sensation and social feeding mediated by a C. elegans guanylate cyclase homologue. Nature. 2004;430(6997):317–322.

12. Zimmer M, Gray JM, Pokala N, et al. Neurons detect increases and decreases in oxygen levels using distinct guanylate cyclases. Neuron. 2009;61(6):865–879.

13. Flibotte JJ, Jablonski AM, Kalb RG. Oxygen sensing neurons and neuropeptides regulate survival after anoxia in developing C. elegans. PLoS One. 2014;9(6):e101102.

14. Van Voorhies WA, Ward S. Broad oxygen tolerance in the nematode Caenorhabditis elegans. J Exp Biol. 2000;203(Pt 16):2467–2478.

15. Padilla PA, Nystul TG, Zager RA, Johnson AC, Roth MB. Dephosphorylation of cell cycle-regulated proteins correlates with anoxia-induced suspended animation in Caenorhabditis elegans. Mol Biol Cell. 2002;13(5):1473–1483.

16. Nystul TG, Goldmark JP, Padilla PA, Roth MB. Suspended animation in C. elegans requires the spindle checkpoint. Science. 2003;302(5647):1038–1041.

17. Pena S, Sherman T, Brookes PS, Nehrke K. The Mitochondrial Unfolded Protein Response Protects against Anoxia in Caenorhabditis elegans. PLoS One. 2016;11(7):e0159989.

18. Doshi S, Price E, Landis J, et al. Neuropeptide signaling regulates the susceptibility of developing C. elegans to anoxia. Free Radic Biol Med. 2019;131:197–208.

19. Mao XR, Crowder CM. Protein misfolding induces hypoxic preconditioning via a subset of the unfolded protein response machinery. Mol Cell Biol. 2010;30(21):5033–5042.

20. Dasgupta N, Patel AM, Scott BA, Crowder CM. Hypoxic preconditioning requires the apoptosis protein CED-4 in C. elegans. Curr Biol. 2007;17(22):1954–1959.

21. Jia B, Crowder CM. Volatile anesthetic preconditioning present in the invertebrate Caenorhabditis elegans. Anesthesiology. 2008;108(3):426–433.

22. Pokala N, Liu Q, Gordus A, Bargmann CI. Inducible and titratable silencing of Caenorhabditis elegans neurons in vivo with histamine-gated chloride channels. Proc Natl Acad Sci U S A. 2014;111 (7):2770–2775.

23. Wang H, Liu J, Gharib S, et al. cGAL’ a temperature-robust GAL4-UAS system for Caenorhabditis elegans. Nat Methods. 2017;14(2):145–148.

24. Stiernagle T. Maintenance of C. elegans. WormBook. 2006:1–11.

25. Mendenhall AR, LaRue B, Padilla PA. Glyceraldehyde-3-phosphate dehydrogenase mediates anoxia response and survival in Caenorhabditis elegans. Genetics. 2006;174(3):1173–1187.

26. Churgin MA, Jung SK, Yu CC, Chen X, Raizen DM, Fang-Yen C. Longitudinal imaging of Caenorhabditis elegans in a microfabricated device reveals variation in behavioral decline during aging. Elife. 2017;6.

27. Rechavi O, Houri-Ze’evi L, Anava S, et al. Starvation-induced transgenerational inheritance of small RNAs in C. elegans. Cell. 2014;158(2):277–287.

28. Artyukhin AB, Yim JJ, Cheong Cheong M, Avery L. Starvation-induced collective behavior in C. elegans. Sci Rep. 2015;5:10647.

29. Cui M, Wang Y, Cavaleri J, Kelson T, Teng Y, Han M. Starvation-Induced Stress Response Is Critically Impacted by Ceramide Levels in Caenorhabditis elegans. Genetics. 2017;205(2):775–785.

30. Ardizzi JP, Epstein HF. Immunochemical localization of myosin heavy chain isoforms and paramyosin in developmentally and structurally diverse muscle cell types of the nematode Caenorhabditis elegans. J Cell Biol. 1987;105(6 Pt 1):2763–2770.

31. Miller DM, 3rd, Ortiz I, Berliner GC, Epstein HF. Differential localization of two myosins within nematode thick filaments. Cell. 1983;34(2):477–490.

32. Schwarz V, Pan J, Voltmer-Irsch S, Hutter H. IgCAMs redundantly control axon navigation in Caenorhabditis elegans. Neural Dev. 2009;4:13.

33. White JG, Southgate E, Thomson JN, Brenner S. The structure of the nervous system of the nematode Caenorhabditis elegans. Philos Trans R Soc Lond B Biol Sci. 1986;314(1165):1–340.

34. Duerr JS, Gaskin J, Rand JB. Identified neurons in C. elegans coexpress vesicular transporters for acetylcholine and monoamines. Am J Physiol Cell Physiol. 2001;280(6):C1616–1622.

35. Chalfie M, Sulston JE, White JG, Southgate E, Thomson JN, Brenner S. The neural circuit for touch sensitivity in Caenorhabditis elegans. J Neurosci. 1985;5(4):956–964.

36. Wicks SR, Rankin CH. Integration of mechanosensory stimuli in Caenorhabditis elegans. J Neurosci. 1995;15(3 Pt 2):2434–2444.

37. Wicks SR, Roehrig CJ, Rankin CH. A dynamic network simulation of the nematode tap withdrawal circuit: predictions concerning synaptic function using behavioral criteria. J Neurosci. 1996;16(12):4017–4031.

38. Lickteig KM, Duerr JS, Frisby DL, Hall DH, Rand JB, Miller DM, 3rd. Regulation of neurotransmitter vesicles by the homeodomain protein UNC-4 and its transcriptional corepressor UNC-37/groucho in Caenorhabditis elegans cholinergic motor neurons. J Neurosci. 2001;21(6):2001–2014.

39. Le Page S, Prunier F. Remote ischemic conditioning: Current clinical perspectives. J Cardiol. 2015;66(2):91–96.

40. Basalay MV, Davidson SM, Gourine AV, Yellon DM. Neural mechanisms in remote ischaemic conditioning in the heart and brain: mechanistic and translational aspects. Basic Res Cardiol. 2018;113(4):25.

41. Marongiu E, Crisafulli A. Cardioprotection acquired through exercise: the role of ischemic preconditioning. Curr Cardiol Rev. 2014;10(4):336–348.

42. Bazopoulou D, Knoefler D, Zheng Y, et al. Developmental ROS individualizes organismal stress resistance and lifespan. Nature. 2019;576(7786):301–305.

43. Herndon LA, Schmeissner PJ, Dudaronek JM, et al. Stochastic and genetic factors influence tissue-specific decline in ageing C. elegans. Nature. 2002;419(6909):808–814.

44. Lithgow GJ, Driscoll M, Phillips P. A long journey to reproducible results. Nature. 2017;548(7668):387–388.

45. Lucanic M, Plummer WT, Chen E, et al. Impact of genetic background and experimental reproducibility on identifying chemical compounds with robust longevity effects. Nat Commun. 2017;8:14256.

46. Rea SL, Wu D, Cypser JR, Vaupel JW, Johnson TE. A stress-sensitive reporter predicts longevity in isogenic populations of Caenorhabditis elegans. Nat Genet. 2005;37(8):894–898.

47. Kuzuya T, Hoshida S, Yamashita N, et al. Delayed effects of sublethal ischemia on the acquisition of tolerance to ischemia. Circ Res. 1993;72(6):1293–1299.

48. Marber MS, Latchman DS, Walker JM, Yellon DM. Cardiac stress protein elevation 24 hours after brief ischemia or heat stress is associated with resistance to myocardial infarction. Circulation. 1993;88(3):1264–1272.

49. Pagliaro P, Gattullo D, Rastaldo R, Losano G. Ischemic preconditioning: from the first to the second window of protection. Life Sci. 2001;69(1):1–15.

50. Gho BC, Schoemaker RG, van den Doel MA, Duncker DJ, Verdouw PD. Myocardial protection by brief ischemia in noncardiac tissue. Circulation. 1996;94(9):2193–2200.

51. Birnbaum Y, Hale SL, Kloner RA. Ischemic preconditioning at a distance: reduction of myocardial infarct size by partial reduction of blood supply combined with rapid stimulation of the gastrocnemius muscle in the rabbit. Circulation. 1997;96(5):1641–1646.

52. Takaoka A, Nakae I, Mitsunami K, et al. Renal ischemia/reperfusion remotely improves myocardial energy metabolism during myocardial ischemia via adenosine receptors in rabbits: effects of “remote preconditioning”. J Am Coll Cardiol. 1999;33(2):556–564.

53. Liem DA, Verdouw PD, Ploeg H, Kazim S, Duncker DJ. Sites of action of adenosine in interorgan preconditioning of the heart. Am J Physiol Heart Circ Physiol. 2002;283(1):H29–37.

54. Rosenberg JH, Werner JH, Moulton MJ, Agrawal DK. Current Modalities and Mechanisms Underlying Cardioprotection by Ischemic Conditioning. J Cardiovasc Transl Res. 2018;11(4):292–307.

55. Yellon DM, Downey JM. Preconditioning the myocardium: from cellular physiology to clinical cardiology. Physiol Rev. 2003;83(4):1113–1151.

56. Dos Santos P, Kowaltowski AJ, Laclau MN, et al. Mechanisms by which opening the mitochondrial ATP-sensitive K(+) channel protects the ischemic heart. Am J Physiol Heart Circ Physiol. 2002;283(1):H284–295.

57. O’Rourke B. Myocardial K(ATP) channels in preconditioning. Circ Res. 2000;87(10):845–855.

58. Ibanez B, Heusch G, Ovize M, Van de Werf F. Evolving therapies for myocardial ischemia/reperfusion injury. J Am Coll Cardiol. 2015;65(14):1454–1471.

59. LaRue BL, Padilla PA. Environmental and genetic preconditioning for long-term anoxia responses requires AMPK in Caenorhabditis elegans. PLoS One. 2011;6(2):e16790.

60. Iranon NN, Jochim BE, Miller DL. Fasting prevents hypoxia-induced defects of proteostasis in C. elegans. PLoS Genet. 2019;15(6):e1008242.

61. Frakes AE, Metcalf MG, Tronnes SU, et al. Four glial cells regulate ER stress resistance and longevity via neuropeptide signaling in C. elegans. Science. 2020;367(6476):436–440.

62. Prahlad V, Cornelius T, Morimoto RI. Regulation of the cellular heat shock response in Caenorhabditis elegans by thermosensory neurons. Science. 2008;320(5877):811–814.

63. Douglas PM, Baird NA, Simic MS, et al. Heterotypic Signals from Neural HSF-1 Separate Thermotolerance from Longevity. Cell Rep. 2015;12(7):1196–1204.

64. Durieux J, Wolff S, Dillin A. The cell-non-autonomous nature of electron transport chain-mediated longevity. Cell. 2011;144(1):79–91.

65. Zhang Q, Wu X, Chen P, et al. The Mitochondrial Unfolded Protein Response Is Mediated Cell-Non-autonomously by Retromer-Dependent Wnt Signaling. Cell. 2018;174(4):870–883

